# Crosstalk between the aryl hydrocarbon receptor and hypoxia inducible factor 1α pathways impairs downstream dioxin response in human islet models

**DOI:** 10.1101/2024.09.25.615065

**Authors:** Noa Gang, Kyle A. van Allen, William G. Willmore, Francis C. Lynn, Jennifer E. Bruin

## Abstract

The incidence of type 2 diabetes (T2D) is increasing globally at a rate that cannot be explained solely by genetic predisposition, diet, or lifestyle. Epidemiology studies report positive associations between exposure to persistent organic pollutants, such as dioxins, and T2D. We previously showed that 2,3,7,8 tetrachlorodibenzo-*p*-dioxin (TCDD) activates the xenobiotic-sensitive aryl hydrocarbon receptor (AHR) in pancreatic islets. The AHR is known to crosstalk with the hypoxia inducible factor 1α (HIF1α) in hepatocytes but whether this crosstalk occurs in islet cells remains unknown. We assessed AHR-HIF1α pathway crosstalk by treating human donor islets and stem cell-derived islets (SC-islets) with TCDD +/- hypoxia and examined the changes in downstream targets of both AHR (e.g., *CYP1A1*) and HIF1α (e.g., *HMOX1*).

SC-islets showed consistent crosstalk between AHR and HIF1α pathways; co-treatment of SC-islets with TCDD + hypoxia robustly suppressed the magnitude of *CYP1A1* induction compared with TCDD treatment alone. In human islets, only 2 of 6 donors showing suppressed *CYP1A1* induction following TCDD + hypoxia co-treatment. In both SC-islets and human donor islets we observed an unexpected hypoxia-mediated suppression of glucose-6-phosphate catalytic subunit 2 (*G6PC2*) expression.

Our study shows AHR-HIF1α crosstalk occurs in both SC-islets and primary human donor islets, but the response of human islets varied between donors. In both models, the HIF1α pathway dominated over the AHR pathway during TCDD + hypoxia co-treatment. Our study is the first to examine whether AHR-HIF1α crosstalk occurs in islet cells and presents novel data on the impact of hypoxia on *G6PC2* gene expression.

## 1.0 Introduction

Diabetes prevalence is increasing rapidly on a global scale. In 2021, 537 million adults worldwide were estimated to be living with diabetes and incidence rate is projected to increase by 46% by 2045 [1]. Type 2 diabetes (T2D) typically develops later in life through myriad factors causing β-cell death or dysfunction, alongside peripheral insulin resistance [2–4]. Traditional risk factors for T2D include genetics, diet, and lifestyle. However, emerging research implicates exposure to man-made, persistent organic pollutants (POPs) as an additional risk factor for T2D incidence [5–9].

POPs are globally distributed, resulting in ubiquitous human exposure [10]. Dioxins and dioxin-like compounds are a class of POPs characterized by a molecular structure of chlorinated benzene rings and a high affinity for the aryl hydrocarbon receptor (AHR) [11,12]. Several epidemiology studies have reported positive associations between dioxin exposure and T2D incidence in diverse populations [6,13–20]. The most toxic and widely studied dioxin is 2,3,7,8-tetrachlorodibenzo-*p*-dioxin (TCDD). Although the mechanism linking TCDD exposure and T2D incidence remains unclear, our laboratory has shown that TCDD exposure to mice *in vivo* and to mouse or human islets *ex vivo* upregulates cytochrome P450 1A1 (*Cyp1a1/CYP1A1*) expression in islet cells [21–23]. The effects of TCDD on glucose tolerance, plasma insulin levels, and insulin sensitivity were abolished in β-cell specific *Ahr* knock-out mice, implicating the AHR pathway in β-cells in mediating the adverse metabolic effects of TCDD [24].

The AHR is a class I bHLH/PAS protein associated with cellular and xenobiotic metabolism [25–30]. Cytosolic AHR is inactive, but when bound by a ligand AHR translocates to the nucleus where the ligand-AHR complex dimerizes to the class II bHLH/PAS protein, aryl hydrocarbon nucleus translocator (ARNT). The AHR-ARNT complex recognizes and binds to xenobiotic response element (XRE) promoter sequences [31–33], leading to upregulation of CYP1A1 enzymes, which are required for the oxidation and subsequent excretion of xenobiotics [28–30,34]. ARNT is a promiscuous Class II protein that shows multiple protein-protein interactions [35–38], including dimerizing with the Class I bHLH/PAS protein hypoxia inducible factor 1α (HIF1α) [36,37,39]. HIF1α is stabilized during periods of insufficient oxygen availability (i.e., hypoxia, < 5% O_2_). Stable HIF1α protein translocates into the nucleus and heterodimerizes with ARNT. The HIF1α-ARNT transcription complex is recognized by hypoxia response element (HRE) promotor sequences that increase gene expression associated with glucose metabolism, cell survival, vascular tone, and vascular growth [33,40–42]

Competitive binding between AHR and HIF1α for ARNT is known to exist in hepatocytes [43]. Interestingly, in hepatocytes and hepatoma cells exposure to hypoxia or hypoxia-mimetics consistently interferes with expression of AHR gene targets, but exposure to AHR ligands does not consistently interfere with the induction of HIF1α gene targets [33,44–47]. For example, mouse hepatoma cells simultaneously exposed to hypoxia and an AHR ligand showed a 30% reduction in the AHR target *Cyp1a1,* but no change in the HIF1α target *Vegf* compared with cells exposed to normoxia and the AHR ligand [48]. Ibrahim et al. (2020) showed that co-treatment of human donor islets with TCDD + proinflammatory cytokines impaired the induction of *CYP1A1* expression compared to TCDD alone [49]. We hypothesized that this inhibitory crosstalk in human islets is like that seen in hepatocytes [50], where HIF1α pathway activation interferes with the AHR pathway.

Human embryonic stem cell (hESC)-derived islet-like endocrine cells (SC-islets) are a developmental research model with potential for clinical diabetes treatment [51]. In the current study we use a genetically modified hESC line that expresses EGFP under control of the *INSULIN* promotor (INS-2A-EGFP cell line [52]), to generate human SC-islets *in vitro*. Although differentiation protocols are continually improving [51,53–56], SC-islets are not fully mature and do not show robust glucose stimulated insulin secretion (GSIS) comparable with primary islets [57–59]. An advantage of SC-islets compared to primary islets from human organ donors is that SC-islets are environmentally naïve and thus have limited biological variability between batches. In contrast, human donor islets are sourced from a diverse population with a broad range of ages, genetic backgrounds, health, lifestyles, and previous environmental exposures. While donor variability is a challenge of studying human islets, this also serves as an important opportunity to examine the range of responses encountered in human islets.

In the current study, we examine AHR-HIF1α pathway crosstalk in human islet cells by exposing primary donor islets and SC-islets to hypoxia alone, TCDD alone, or hypoxia + TCDD combined for 48 hours. We measured gene targets of AHR (e.g., *CYP1A1*) as an indicator of AHR activation and gene targets of HIF1α (e.g., *HMOX1*) as an indication of HIF1α activation. GSIS was also measured in primary islets to determine the effect of AHR-HIF1α crosstalk on human islet function.

## 2.0 Methods

### 2.1 hESC differentiation and maintenance

SC-islets were derived from INS-2A-EGFP WA01 hESCs by Dr. Francis Lynn (University of British Columbia, BC, Canada) and differentiated over 27-days [55,60]. SC-islets were then shipped from Vancouver to Ottawa and maintained in Stage 6 media [CMRL-1066 (VWR, #CA45001-114; Radnor, PA) containing 20 g/L bovine serum albumin (Sigma-Aldrich #10775835001), 1X Glutamax (Gibco #35050079), 1% penicillin-streptomycin (Gibco #15140122), 0.5 mM pyruvate (Sigma-Aldrich #S8636-100 mL), 0.5X Insulin Transferrin-Selenium-X (Thermo Fisher #51500056), 35 nM zinc sulfate (Sigma-Aldrich #Z0251-100G), 1 mM N-acetyl-L-cysteine (Sigma-Aldrich #A9165-5G), 10 ug/L Heparin (Sigma-Aldrich #H3149-10KU), 1:2000 Trace elements A (Corning 25-021-CI), 1:2000 Trace Elements B (Corning 25-022-CI #MT99175CI), 10 nM T3 (Sigma, #T6397-100MG), 0.5 μM ZM 447439 (Cedarlane Labs, #S1103-10MM/1ML; Burlington, ON), and 1:2000 lipid concentrate (Fisher Scientific #11905031)]. SC-islets were maintained at 37°C, 21% O_2_, and 5% CO_2_ on an orbital shaker (106 RPM) until analyses, with media changes every two days. SC-islets are composed of a mixture of pancreatic endocrine cells, including ∼30-70% INS-GFP+ cells. SC-islets aged to day 50-60 show a modest GSIS response [55]. Therefore, we used SC-islets aged day 50+ in our experiments. Each SC-islet differentiation was considered one “biological replicate” (n=3 technical replicates per condition per differentiation; n=3 biological replicates).

### 2.2 Human organ donor islet cell culture

Human islets were provided from the Alberta Diabetes Institute IsletCore at the University of Alberta (www.bcell.org/adi-isletcore) with the assistance of the Human Organ Procurement and Exchange (HOPE) program, Trillium Gift of Life Network (TGLN), and other Canadian organ procurement organizations. Islet isolation was approved by the Human Research Ethics Board at the University of Alberta (Pro00013094). All donors’ families gave informed consent for the use of pancreatic tissue in research. All research using human islets was approved by the Research Ethics Board at Carleton University. Islet purity ranged between 75-90% for this study (**Table 1**). Donor islets were shipped overnight to Ottawa in CMRL media (Fisher Scientific, #11-530-037). Upon arrival, islets were transferred to low glucose DMEM media (LG-DMEM; Fisher Scientific, #11-885-084) supplemented with 10% FBS (Sigma-Aldrich #F1051-500ML) and 1% penicillin-streptomycin (Gibco, #15140122; Billings, MT) and cultured at 37°C, 21% O_2_, and 5% CO_2_ overnight to allow for recovery. See **Table 1** for donor characteristics. Human islets from separate donors were considered as biological replicates (n=3-5 technical replicates per condition per donor; n=3 biological replicates per sex).

**Table 1.** Human islet donor characteristics.

| Donor | Sex | Age (years) | BMI | HbA1C (%) | Cold ischemia time (hours) | Islet purity (%) |
| --- | --- | --- | --- | --- | --- | --- |
| R425 | Female | 36 | 25.2 | 5.3 | 12 | 80 |
| R427 | Male | 52 | 35.8 | 5.8 | 10 | 90 |
| R430 | Male | 49 | 24.4 | 4.8 | 8 | 90 |
| R434 | Female | 50 | 24.2 | 5.7 | 5 | 85 |
| R444 | Female | 48 | 22.0 | 5.3 | 9 | 80 |
| R461 | Male | 61 | 31.6 | 4.3 | 21 | 75 |

### 2.3 Experimental conditions

Two experimental paradigms were used in this study (**Figure 1**). First, we treated SC-islets with TCDD +/- hypoxia exposure continuously for 48 hours or pre-treated SC-islets with either TCDD alone *or* hypoxia alone for 24 hours prior to co-treatment with TCDD + hypoxia for 24 hours (**Figure 1A**). Lastly, we compared the effects of TCDD +/- hypoxia co-treatment for 48 hours in SC-islets versus human donor islets (**Figure 1B**).

**Figure 1.**
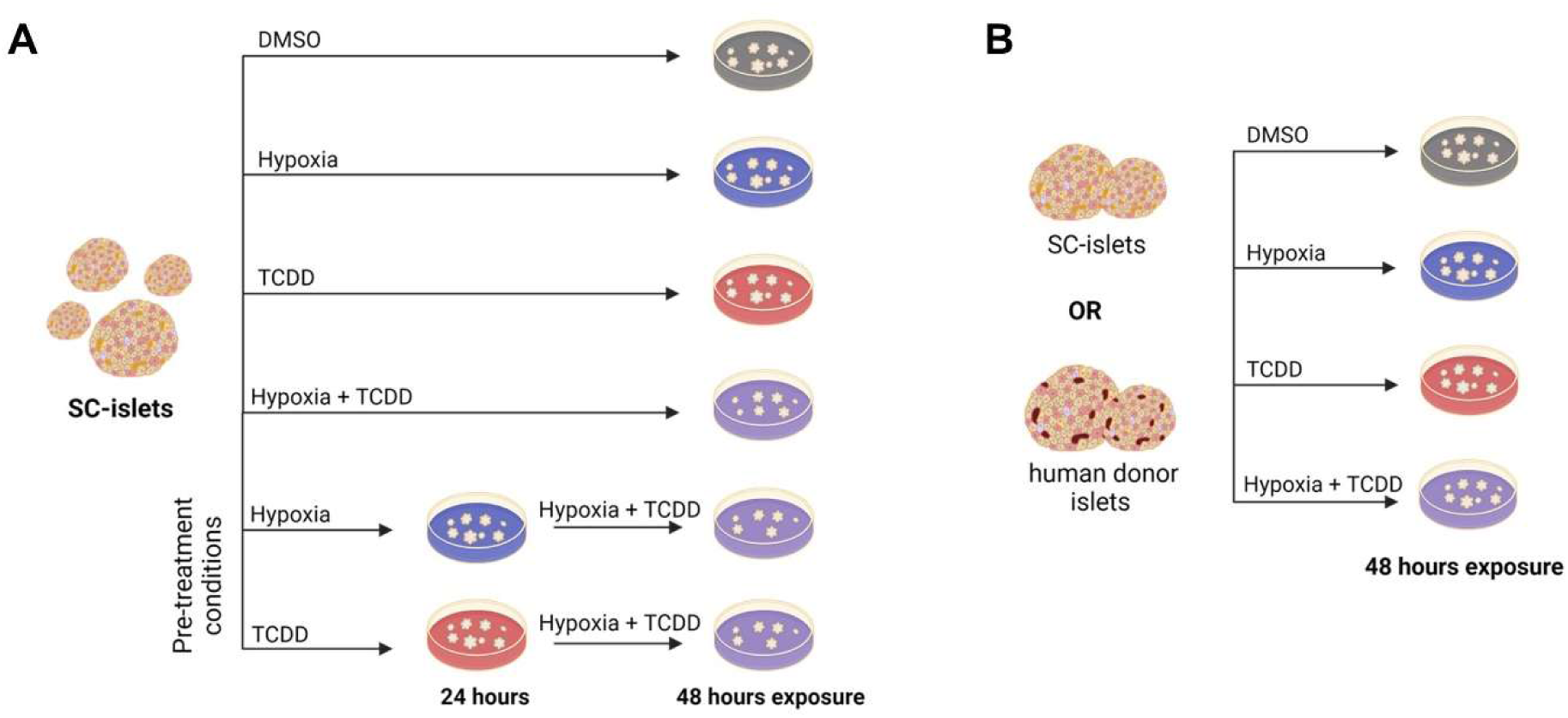
Summary of treatment conditions applied to SC-islets and human donor islets. **(A)** SC-islets were exposed to TCDD (10 nM) +/- hypoxia (1% O_2_) for 48 hours continuously or to TCDD or hypoxia pre-treatment for 24 hours followed by TCDD + hypoxia co-treatment for 24 hours. **(B)** SC-islets or human donor islets were exposed to TCDD +/- hypoxia co-treatment continuously for 48 hours. Made with BioRender.com.

SC-islets or human donor islets were hand-picked and transferred to 6-well non-TC treated plates (VWR, #10861-554) to undergo the treatment condition outlined above: (i) DMSO (#276855-100ML, Sigma-Aldrich, Oakville, ON, Canada) vehicle control; (ii) normoxia (21% O_2_, 5% CO_2_); (iii) 10 nM TCDD (#AD404SDMSO10X1ML, Accustandard, Chromatographic Specialties, Brockville, ON, Canada); (iv) hypoxia (1% O_2_) in a triple-gas (N_2_-O_2_-CO_2_) incubator (ThermoForma, Series II, 3130, Marietta, Ohio).

### 2.4 Quantitative real time PCR

Immediately following treatment, SC-islets or human islets were transferred to buffer RLT (#79216, Qiagen, Hilden, Germany) and stored at -80°C. mRNA was isolated using RNeasy Micro kit (#74004, Qiagen, Hilden, Germany) as per the manufacturer’s instructions. DNase treatment was performed prior to cDNA synthesis with the iScriptTM gDNA Clear cDNA Synthesis Kit (#1725035, Bio-Rad, Mississauga, ON, Canada). Real time (RT) qPCR was performed using SsoAdvanced Universal SYBR Green Supermix (Bio-Rad, #1725271) and run on a CFX394 (Bio-Rad). *PPIA* was used as the reference gene. Primer sequences are listed in **Table S1**. Data were analyzed using the 2ΔΔCT relative quantitation method.

### 2.5 Glucose-stimulated insulin secretion

SC-islets have limited glucose responsiveness in our hands, therefore only human islets were used to measure GSIS (n=3-5 technical replicates per donor; n=6 biological replicates, i.e., donors). Immediately following treatment, 20 islets per condition were moved to pre-warmed (37°C) Krebs–Ringer bicarbonate buffer (KRBB) solution (115 mmol/l NaCl, 5 mmol/l KCl, 24 mmol/l NaHCO_3_, 2.5 mmol/L CaCl_2_, 1 mmol/l MgCl_2_, 10 mmol/l HEPES, 0.1% (wt/vol.) BSA, pH 7.4) with 2.8 mmol/L glucose (low glucose, LG) for a 60 min pre-incubation in 1.5 mL Eppendorf tubes. Supernatant was removed and replaced with 500 μL of LG KRBB for 1 h, followed by transfer to 500 μL of KRBB with 16.7 mmol/L glucose (high glucose, HG) for 1 h at 37°C. The LG KRBB and HG KRBB samples were centrifuged, and the supernatant stored at −20°C. To measure insulin content, supernatant was removed and replaced with an acid-ethanol solution of 1.5% vol/vol HCl in 75% vol/vol ethanol at 4°C overnight and neutralised with 1 mol/L Tris base (pH 7.5) before long-term storage at −20°C. Concentrations of human insulin were measured by ELISA (#80-INSHU-CH10, ALPCO, Salem, NH, USA).

### 2.6 Statistics

All statistics were performed using GraphPad Prism 10.0.0 (GraphPad Software Inc., La Jolla, CA). For all analyses, p<0.05 was considered statistically significant and any potential outliers were tested for significant using a Grubbs’ test with α=0.05. Variance between biological replicates was tested for using a nested one-way ANOVA. Non-parametric statistics were used in cases where the data failed normality or equal variance tests. Data are presented as means ± SEM. Figure legends specify whether individual datapoints are technical or biological replicates.

### 3.0 Results

### 3.1 Co-exposure of SC-islets to TCDD + hypoxia impairs CYP1A1 induction by TCDD, regardless of pre-treatment conditions

To determine if hypoxia interferes with TCDD-mediated AHR activation in islets, we measured *CYP1A1* and *HMOX1* induction in SC-islets following exposure to TCDD +/- hypoxia (Figure 1). As expected, TCDD significantly upregulated *CYP1A1* expression in SC-islets ∼41-fold compared to vehicle (**Figure 2A**). However, when SC-islets were co-treated with TCDD + hypoxia, the magnitude of *CYP1A1* induction was suppressed to ∼21-fold (**Figure 2A**). Hypoxia exposure induced *HMOX1* ∼24-fold compared to normoxia in SC-islets and this was not affected by co-treatment with TCDD (**Figure 2B**).

**Figure 2.**
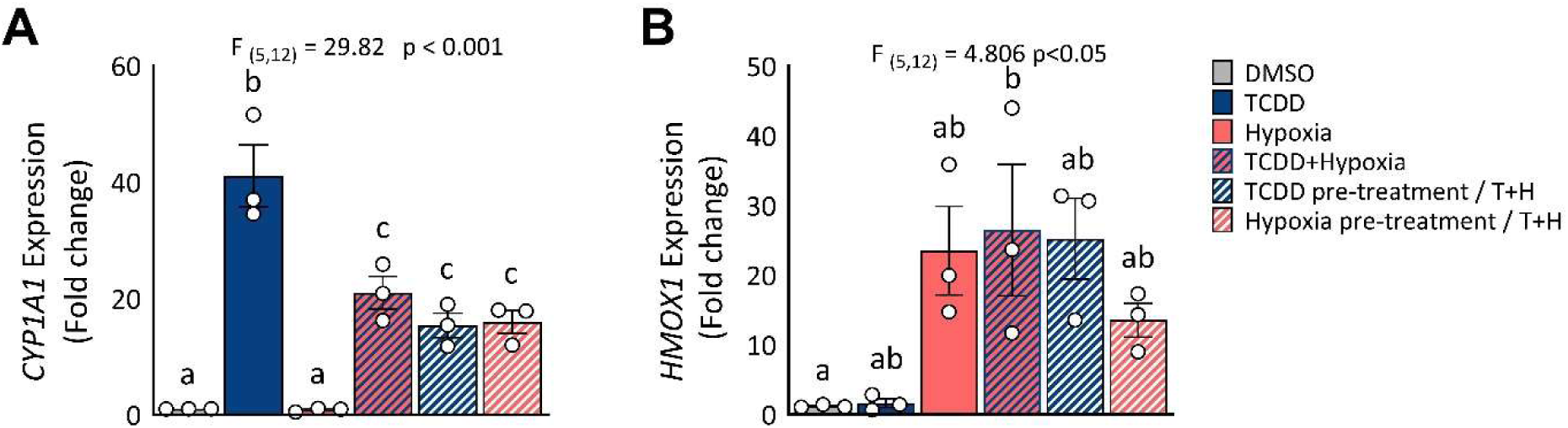
Hypoxia interfered with *CYP1A1* upregulation by TCDD in SC-islets. **(A)** *CYP1A1* and **(B)** *HMOX1* expression in SC-islets co-treated with TCDD +/- hypoxia for 48 hours or following either TCDD or hypoxia pre-treatment for 24 hours and TCDD + hypoxia co-treatment for 24 hours. Gene expression is relative to DMSO-treated SC-islets. Data are mean +/- SEM and individual data points represent biological replicates (i.e. different differentiations). Significance was determined by a one-way ANOVA and Tukey post-hoc test. Different letters indicate statistically significant differences (p<0.05).

We also investigated whether the degree of crosstalk between AHR and HIF1α in SC-islets is influenced by which pathway is activated first—i.e., a first-come, first-serve effect. We hypothesized that activation of either the AHR or HIF1α pathway *before* a competitive co-exposure condition would dictate the dimerization partner of ARNT. SC-islets were pre-treated with either TCDD or hypoxia for 24 hours prior to co-treatment with TCDD + hypoxia for 24 hours (**Figure 1A**). We found that all TCDD + hypoxia co-treatment conditions significantly impaired the degree of *CYP1A1* induction compared to treatment with TCDD alone (**Figure 2A**). Pre-treatment of SC-islets with either TCDD or hypoxia did not alter this effect (**Figure 2A**). Similarly, we found that all hypoxia treatment conditions significantly upregulated *HMOX1* in SC-islets, irrespective of co-treatment or pre-treatment with TCDD (**Figure 2B**).

### 3.2 Co-exposure of SC-islets to TCDD + hypoxia also causes an interaction effect on G6PC2

To further explore the impact of molecular crosstalk in SC-islets, we focused on the first 4 conditions: vehicle, TCDD alone, hypoxia alone, or TCDD + hypoxia for 48-hours (**Figure 1B**). We measured *CYP1A1* expression alongside other AHR targets (*AHR*, *AHRR*), HIF1α targets (*HMOX1*, *VEGFA)*, *ARNT,* biomarkers of β-cell function/identity (*MAFA, SLC2A1, G6PC2*), and a marker of oxidative stress (*MNSOD)* (**Figure 3**). As shown in **Figure 2A** and re-plotted here, there was a significant interaction effect of TCDD + hypoxia co-treatment on *CYP1A1* expression in SC-islets (**Figure 3A**). The AHR repressor gene, *AHRR,* was significantly upregulated by TCDD exposure, as expected, but unlike *CYP1A1* there was no interaction effect with hypoxia (**Figure 3C**). Exposure of SC-islets to hypoxia significantly upregulated *AHR*, *HMOX1, VEGFA, MNSOD,* and *SLC2A1* (**Figure 3B,D,E,G,H**) and downregulated *MAFA* expression (**Figure 3I**); none of these changes were affected by co-treatment with TCDD. Both TCDD and hypoxia exposure alone modestly upregulated *ARNT* expression, but there was no interaction effect on *ARNT* (**Figure 3F**). Interestingly, an unexpected interaction effect was seen for *G6PC2* expression (**Figure 3J**). Both TCDD alone and hypoxia alone significantly downregulated *G6PC2* expression compared with vehicle/normoxia control conditions (**Figure 3J**). Hypoxia-mediated downregulation dominated the interaction effect seen following TCDD + hypoxia co-treatment (**Figure 3J**).

**Figure 3.**
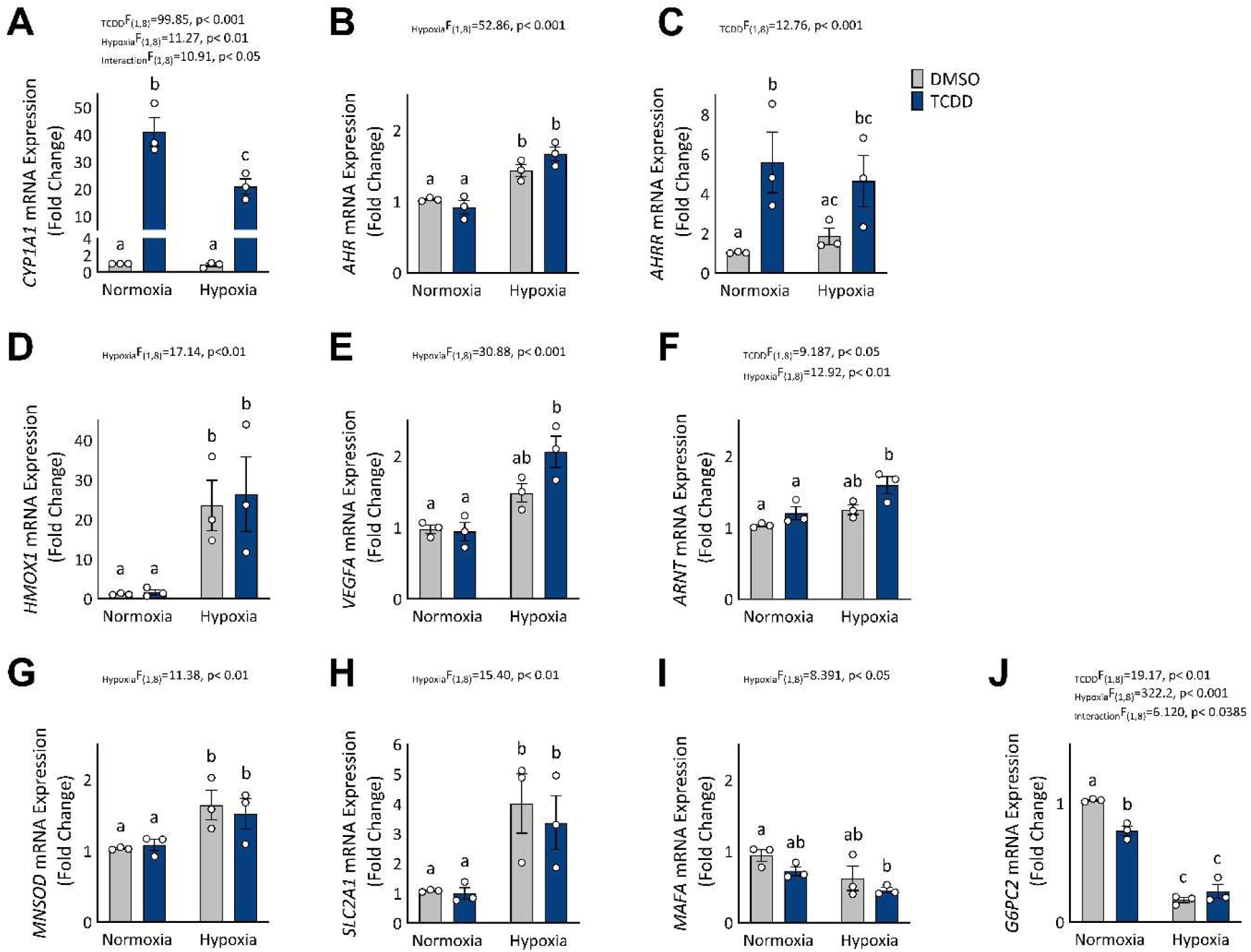
TCDD + hypoxia co-treatment causes interactive effect for *CYP1A1* and *G6PC2* expression in SC-islets. SC-islets were exposed to TCDD +/- hypoxia for 48 hours and gene expression was measured for (A) *CYP1A1*, (B) *AHR*, (C) *AHRR*, (D) *HMOX1*, (E) *VEGFA*, (F) *ARNT*, (G) *MNSOD*, (H) *MAFA*, (I) *SLC2A1*, and (J) *G6PC2*. Gene expression is relative to DMSO-treated SC-islets. Data are mean +/- SEM and individual data points represent biological replicates (i.e. different differentiations). Significance was determined by a two-way ANOVA and Tukey post-hoc test. Different letters indicate statistically significant differences (p<0.05).

### 3.3 TCDD consistently induces CYP1A1 in human donor islets but AHR-HIF1α crosstalk varies between donors

We next examined whether the crosstalk between AHR and HIF1α pathways also occurs in primary human islets from deceased organ donors. Human islets from 3 male donors and 3 female donors were included in our study. Average donor age was 49.3 (+/- 8.0) years, BMI was 27.2 (+/- 5.3), and HbA1c was 5.2 (+/- 0.6) (**Table 1**). Human donor islets were co-exposed to TCDD +/- hypoxia continuously for 48 hours, as described for the SC-islets (**Figure 1B**). Unlike SC-islets, human islets showed significant variability between biological replicates for the gene targets examined (Χ^2^ =26.17-239.4, all p<0.0001). These variable effects are summarized in **Table 2**, detailed statistics are provided in **Supplementary Table 2**, and additional donor information can be found at humanislets.com [61].

**Table 2.** SC-islets and human donor islets treated with TCDD +/- hypoxia show variability for gene expression changes. A summary table of the main effects of TCDD +/- hypoxia on SC-islets and human donor islets for expression of *CYP1A1, HMOX1, AHRR, ARNT,* and *G6PC2.* Bolded fold-increase indicates significance for TCDD or hypoxia treatments alone, compared with vehicle control.

| Cell source | *CYP1A1* |  |  | *HMOX1* |  |  | *AHRR* |  |  | *ARNT* |  |  | *G6PC2* |  |  |
| --- | --- | --- | --- | --- | --- | --- | --- | --- | --- | --- | --- | --- | --- | --- | --- |
|  | TCDD | Hypoxia | Crosstalk | TCDD | Hypoxia | Crosstalk | TCDD | Hypoxia | Crosstalk | TCDD | Hypoxia | Crosstalk | TCDD | Hypoxia | Crosstalk |
| SC-Islets | ↑ (41-fold) | ↑ (1.1-fold) | ✓ | - | ↑ (24-fold) | X | ↑ (6-fold) | - | X | ↑ (1.2-fold) | ↑ (1.2-fold) | X | ↓ | ↓ | ✓ |
| R425-♂ | ↑ (10-fold) | ↓ | ✓ | - | ↑ (2-fold) | ✓ | ↑ (1.3-fold) | - | ✓ | - | - | X | - | ↓ | X |
| R434-♂ | ↑ (3-fold) | - | X | ↑ (1.4-fold) | ↑ (5-fold) | X | ↑ (2-fold) | - | X | ↑ (1.7-fold) | ↑ (1.7-fold) | X | ↑ (2-fold) | ↓ | X |
| R444-♂ | ↑ (11-fold) | - | X | ↑ (3-fold) | ↑ (3-fold) | X | ↑ (3-fold) | - | X | ↑ (2-fold) | - | X | ↑ (2-fold) | - | X |
| R427-♀ | ↑ (3-fold) | ↓ | X | - | ↑ (3-fold) | X | - | - | X | - | - | X | - | ↓ | X |
| R430-♀ | ↑ (3-fold) | - | ✓ | - | ↑ (3-fold) | X | ↑ (1.1-fold) | - | ✓ | - | - | ✓ | - | ↓ | X |
| R461-♀ | ↑ (15-fold) | ↑ (2-fold) | ✓ | - | ↑ (3-fold) | X | ↑ (2-fold) | - | X | - | - | X | - | ↓ | X |

TCDD exposure significantly increased *CYP1A1* expression in all donors, albeit to varying degrees (2.5-fold to 15.8-fold, **Figure 4A**, **Table 2**). We saw a significant interaction effect for TCDD + hypoxia co-treatment to suppress the magnitude of *CYP1A1* induction compared with TCDD alone in 2 of the 6 donors (Donors R425 and R461, **Figure 4A**, **Table 2**). There was also a trending interaction effect in Donor R430 (p = 0.0552, **Figure 4A**, **Table 2**). No interaction effect for *CYP1A1* expression was found in the other donor islets co-exposed to TCDD + hypoxia (**Figure 4A**, **Table 2**).

**Figure 4.**
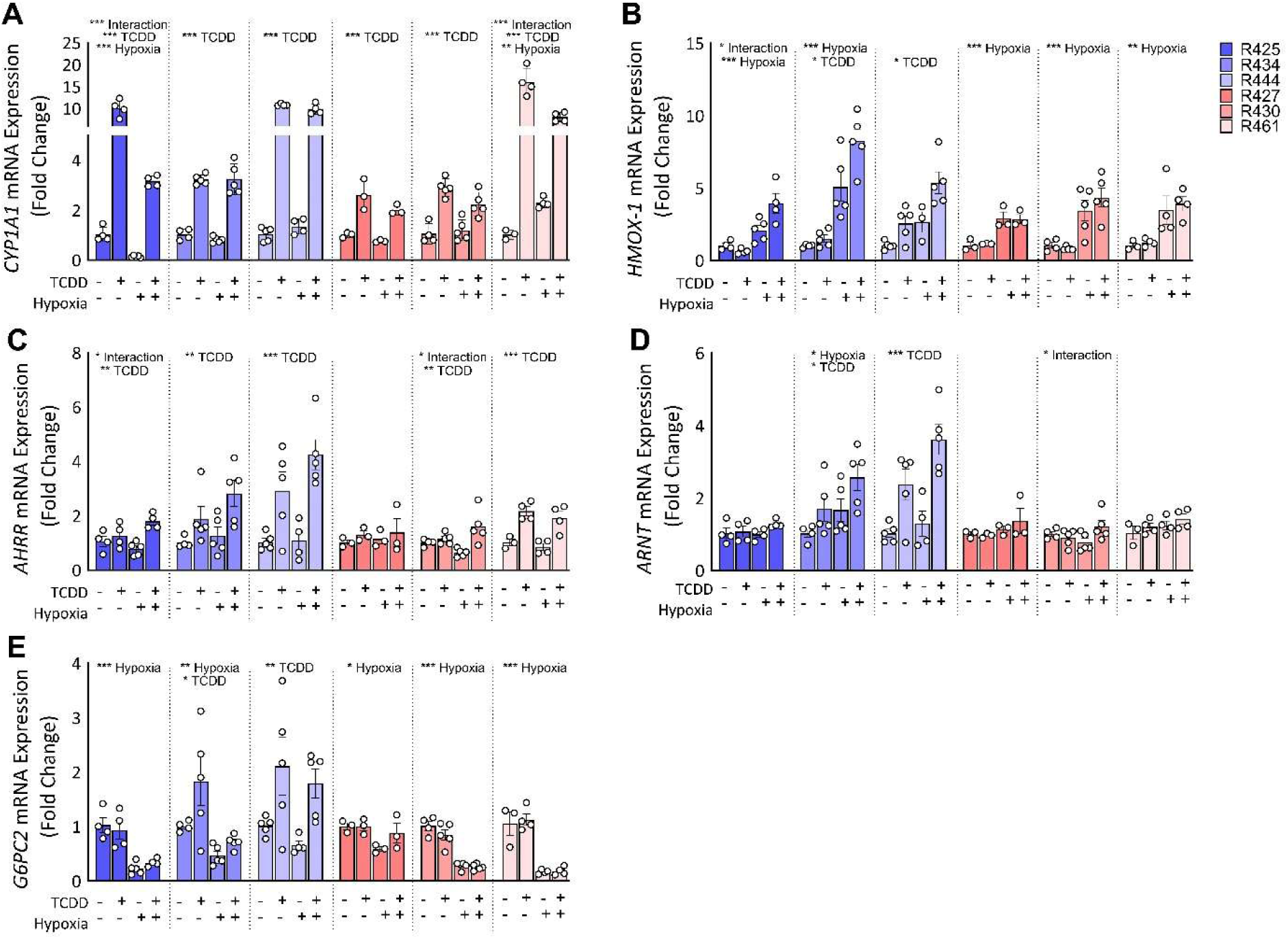
Co-treatment of human islets to TCDD + hypoxia impairs induction of *CYP1A1* by TCDD in 2 of 6 donors. Human islets were exposed to TCDD +/- hypoxia for 48 hours and gene expression was measured for (A) *CYP1A1,* (B) *HMOX1,* (C) *AHRR,* (D) *ARNT,* and (E) *G6PC2.* Gene expression is relative to DMSO-treated islets. Female donors are indicated in blue and male donors are indicated in red. Data are mean +/- SEM and individual data points represent technical replicates. Significance was determined by a two-way ANOVA and Dunnett post-doc test. Statistically significant differences within donors are indicated by an asterisk (*p<0.05, **p<0.01, ***p<0.001).

### 3.4 TCDD consistently upregulates AHRR expression in human islets; hypoxia consistently induces HMOX1 and downregulates G6PC2 expression in human islets

All donors showed a main effect or trend for modest *HMOX1* upregulation by hypoxia (2.2-fold to 5.0-fold, **Figure 4B**, **Table 2**). A modest interaction effect by TCDD + hypoxia co-exposure augmenting *HMOX1* upregulation was seen for one donor (R425, **Figure 4B**, **Table 2**).

All donors except one (Donor R427) showed a main effect for TCDD exposure upregulating *AHRR* expression (**Figure 4C**, **Table 2**); this is consistent with the effect seen in SC-islets (**Figure 3C**, **Table 2**). In addition, two donors (R425 and R430) showed a modest TCDD + hypoxia interaction effect for *AHRR* (**Figure 4C**, **Table 2**).

Changes in *ARNT* expression were highly variable between donors. Three donors showed no change in *ARNT* expression across treatments (Donors R425, R427, R461; **Figure 4D**, **Table 2**). Donor R430 showed a modest interaction effect, Donor R434 showed a main effect for both hypoxia and TCDD, and Donor R444 showed a main effect for TCDD exposure only (**Figure 4D**, **Table 2)**.

All human donors except R444 showed a significant main effect for hypoxia to downregulate *G6PC2* expression (**Figure 4E**, **Table 2**). Donors R434 and R444 additionally showed a main effect for *G6PC2* upregulation by TCDD (**Figure 4E**, **Table 2**). TCDD-driven upregulation of *G6PC2* in donor islets contrasts the modest but consistent TCDD-driven *G6PC2* downregulation seen in the SC-islets (**Figure 3J**, **Table 2**). There were no interaction effects found for *G6PC2* expression among the donor islets, which also differs from the significant interaction effect seen in SC-islets.

### 3.5 Hypoxia, but not TCDD, negatively impacts donor islet GSIS

To determine whether TCDD +/- hypoxia exposure impacted islet function we conducted a GSIS assessment immediately following 48-hour exposure conditions. Across all donors, TCDD did not impact basal or glucose-stimulated insulin secretion compared with vehicle control (**Figure 5A**,**B**). Exposure to hypoxia dramatically reduced insulin secretion in response to high glucose (**Figure 5A**,**B**). This effect was maintained but not amplified when cells were co-treated with hypoxia + TCDD (**Figure 5A**,**B**). Hypoxia treatment regardless of co-treatment with TCDD resulted in mild reduction of human islet total insulin content (**Figure 5C**).

**Figure 5.**
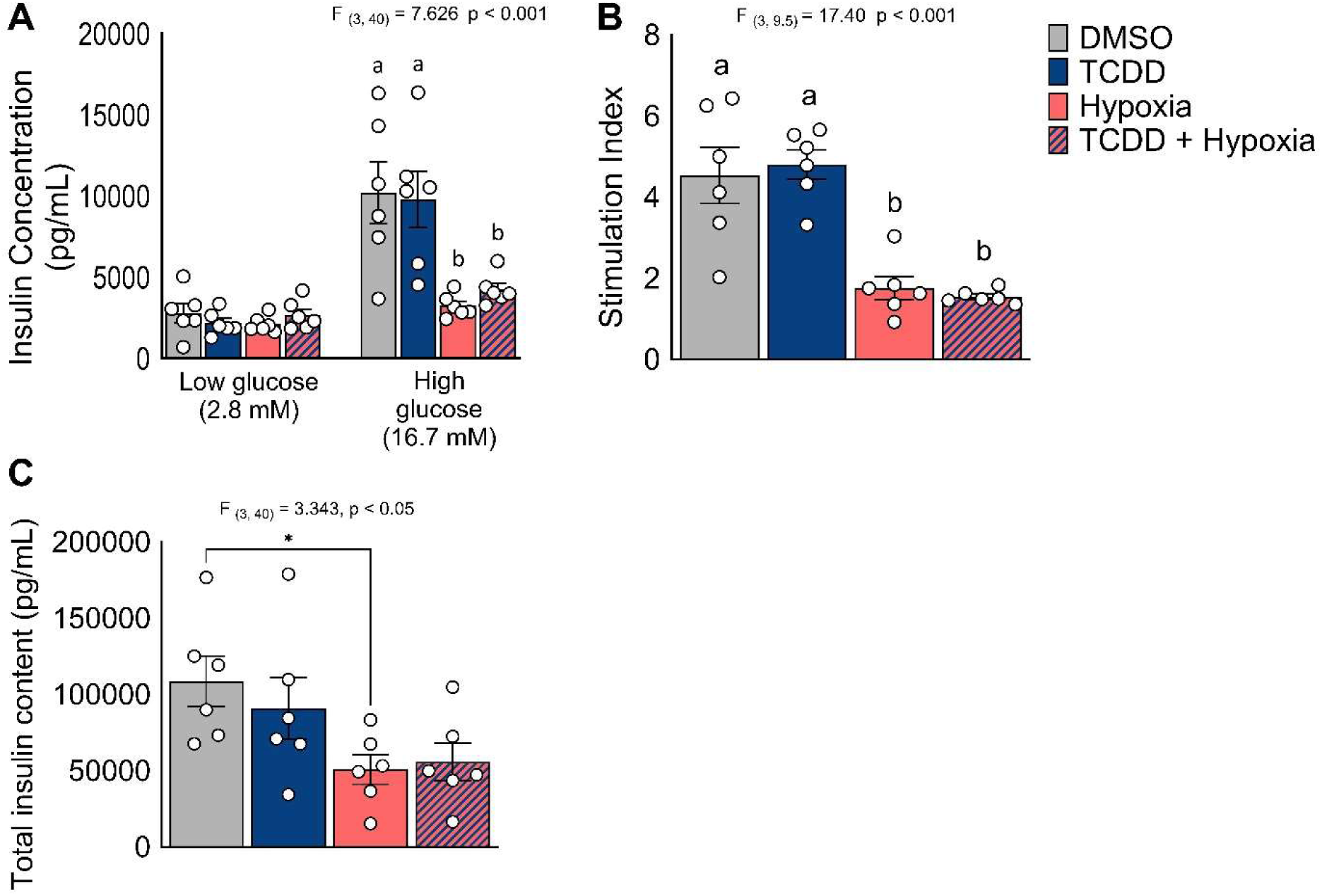
Hypoxia exposure negatively impacts glucose stimulated insulin secretion. Human islets were exposed to TCDD +/- hypoxia for 48 hours and **(A)** insulin secretion was measured following 1 hour in low (2.8 mM) and high (16.7 mM) glucose concentrations. **(B)** Stimulation index is expressed as secreted insulin concentration following high glucose relative to low glucose stimulation. **(C)** Total insulin content following acid ethanol lysis. Data are mean +/- SEM and individual data points represent biological replicates (i.e. different islet donors). Significance was determined with **(A)** a two-way ANOVA and Šidák post-hoc test, **(B)** a Brown-Forsythe ANOVA and Dunnett post-hoc test, and **(C)** a one-way ANOVA and Dunnett post-hoc test. Different letters and asterisks indicate statistically significant differences (*p<0.05).

### 4.0 Discussion

This study investigated whether there is crosstalk between AHR and HIF1α pathways in SC-islets and human donor islets. TCDD exposure consistently activated the AHR pathway, as indicated by *CYP1A1* upregulation, in SC-islets and all human islet donors. However, the degree of *CYP1A1* induction was more pronounced in SC-islets than human islets and varied substantially between human islet donors [62], [63]. Hypoxia exposure consistently interfered with TCDD-mediated *CYP1A1* induction in the SC-islet model, a clear indication of AHR-HIF1α crosstalk. We also saw evidence of AHR-HIF1α crosstalk in human islets but only in 2 of 6 donors.

There are numerous reasons why AHR-HIF1α crosstalk may be more consistent in SC-islets than human donor islets. First, we speculate that previous environmental exposures likely influence the sensitivity of donor islets to TCDD and/or hypoxia exposure *ex vivo*. SC-islets are environmentally naive whereas human islets have been exposed to a broad variety of environmental contaminants throughout their lifetime [64–70], which impact baseline AHR pathway activation between donors. Furthermore, dietary habits, smoking or drug use, and other environmental factors can lead to cellular hypoxia [71–73], which likely impacts baseline HIF1α pathway activation. Donor age is another important source of biological variation; the human islets used in our study were isolated from donors ranging between 36 to 61 years of age. AHR activation and downstream effects are known to vary with age [74,75] and baseline *AHR* and *ARNT* levels vary by age and tissue state [76]. Lastly, the diverse genetic background of islet donors likely contributed to variable responses compared to the uniform genetic background of SC-islets derived from a single donor. In summary, the differences in robustness and variability of AHR-HIF1α crosstalk between SC-islets and human donor islets likely arises from a mixture of these considerations.

Our study demonstrated a clear dominance of the HIF1α pathway over the AHR pathway in islets, even when cells were pre-treated with TCDD for 24 hours. This was evident in the lack of interference by TCDD with hypoxia-mediated *HMOX1* induction in both SC-islets and human islets. Notably, there is little to no evidence of ARNT-independent gene induction by HIF1α [77–81]. The dominance of hypoxia-mediated effects was also clear in the GSIS response of human islets. Hypoxia exposure profoundly blunted the GSIS response, as expected based on previous findings [82–87], and this effect was seen irrespective of co-treatment with TCDD. Our data suggest that ARNT binding affinity is critical to AHR-HIF1α crosstalk in human islets. Gradin *et al.* used coimmunoprecipitation to show stabilized HIF1α has a high binding affinity with ARNT in a human hepatoma cell line. Furthermore, this group found the HIF1α pathway dominates over AHR in binding with ARNT in a rabbit reticulocyte lysate [50]. The exact mechanism(s) driving the HIF1α-ARNT binding dominance over AHR-ARNT in SC-islets and human donor islets warrants further investigation.

The presence of AHR-HIF1α crosstalk in SC-islets and human donor islets has interesting implications for islet transplantation. Human islets are transiently hypoxic following clinical transplantation [88,89] and need to mount an efficient and robust HIF1α response to induce vascularization. Given that the AHR and HIF1α pathways both rely on access to ARNT, it is possible that sustained exposure to an AHR ligand(s) could interfere with the HIF1α-mediated hypoxia response post-transplant of either SC-islets or human donor islets. While our experiment models showed the HIF1α pathway dominating AHR-HIF1α crosstalk in islets, this was a short-term (48-hr) window of crosstalk *in vitro*. It remains possible that long-term exposure of patients to AHR ligands post-transplant could compromise engraftment via interfering with the HIF1α pathway. Given that we do not fully understand the role of AHR pathway activation within islets, it is also difficult to appreciate the potential impact that post-transplant hypoxia may have on islet health if HIF1α activation interferes with the AHR-mediated detoxification response in engrafted islets.

The hypoxia-mediated downregulation of *G6PC2* was an unexpected finding in both SC-islets and human donor islets. The consistency of this effect between donors is particularly notable. The consistent changes in *HMOX1, CYP1A1,* and *AHRR* expression were expected based on well-established effects of hypoxia [41,88] and TCDD [37], in islets and other cell types. Hypoxia exposure upregulates *G6pc* expression in mouse liver tissue and in a human liver cell line (HepG2) [90,91], but to our knowledge the connection between hypoxia-mediated *G6PC2* downregulation in human islets has not been previously described. In cancer patients and cancer cell lines *G6PC2* was described as part of the battery of gene changes associated with the Warburg effect (aerobic glycolysis) [92,93], and hypoxia-exposed rats showed *G6PC2* downregulation in liver tissue [94]. In humans the *G6PC2* isoform is islet-specific [95], associated with autoimmunity [96], and responsible for tight regulation of fasting blood glucose levels [97–100]. *G6PC2* expression is downregulated in islets from donors diagnosed with T2D, a condition associated with islet hypoxia [86,101–103]. We speculate that beyond regulating blood glucose levels in diabetes, downregulation of *G6PC2* is protective by increasing intracellular glucose levels available for anaerobic glycolysis and ATP production necessary for insulin secretion. In support, Rahim et al. (2022) showed *G6pc2* knock-out improved GSIS response by modulating glycolysis in a mouse pancreatic β-cell line [104]. Hypoxia-mediated *G6PC2* downregulation may work in concert with other hypoxia-mediated gene expression changes in the islet known to promote glycolysis and suppress gluconeogenesis [88,105].

### 5.0 Conclusions

In conclusion, our study is the first to show evidence for AHR-HIF1α crosstalk in islets. This crosstalk was dominated by the HIF1α pathway and was consistently observed in SC-islets. However, in human islets, the extent to which *CYP1A1* expression was upregulated following TCDD exposure and the presence of AHR-HIF1α crosstalk varied substantially between donors. We believe further study is warranted to examine why HIF1α dominates AHR-HIF1α crosstalk in islets, and what factors contribute to the variability seen in human donor islets. Furthermore, we believe the previously unreported finding of hypoxia-mediated *G6PC2* downregulation in SC-islets and human donor islets requires further investigation.

## Acknowledgements

Human islets for research were provided by the Alberta Diabetes Institute IsletCore at the University of Alberta in Edmonton (http://www.bcell.org/adi-isletcore.html) with the assistance of the Human Organ Procurement and Exchange (HOPE) program, Trillium Gift of Life Network (TGLN), and other Canadian organ procurement organizations. We thank the IsletCore staff, the islet donors, and their families.

We thank Dr. Jan Mennigan for thoughtful discussions over experimental design. We thank Dr. Ji Soo Yoon and Ekaterina Filatov for their valuable help designing figures.

The authors acknowledge that Carleton University is situated on the traditional, ancestral, and unceded territories of the Nehiyahwak, Nahikawe, Dakata, Lakota, Dene, and Metis Nations. The authors acknowledge that UBC and BC Children’s Hospital are situated on the traditional, ancestral, and unceded territories of the Coast Salish peoples—the Sḵwxwú7mesh (Squamish), Səlı’lwətaʔ/Selilwitulh (Tsleil-Waututh), and xʷməqkʷəyəm (Musqueam) Nations.

## Author Contributions

**NG:** Conceptualization, Methodology, Formal analysis, Investigation, Writing – Original draft, Writing – Review & editing, Visualization, Funding acquisition. **JEB:** Conceptualization, Methodology, Formal analysis, Resources, Writing – Review & editing, Visualization, Supervision, Funding acquisition. **WGW:** Conceptualization, Methodology, Resources, Writing – Review & editing. **KVA:** Investigation, Writing – Review & editing. **FCL:** Investigation, Resources, Writing – Review & editing.

## Funding

This research was supported by a Canadian Institutes of Health Research-Juvenile Diabetes Research Federation (CIHR-JDRF) Team Grant (#ASD-173663/5-SRA-2020-1059-S-B) and Natural Sciences and Engineering Research Council (NSERC) Discovery Grants to JEB (#RGPIN-2017-06265). NG was supported by an NSERC Canadian Graduate Student-Doctoral (CGS-D) award and KAV was supported by an NSERC-Collaborative Research and Training Experience (CREATE) award. JEB is supported by an Early Researcher Award from the Ontario Government.

## Abbreviations

AHR: Aryl hydrocarbon receptor
AHRR: Aryl hydrocarbon receptor repressor
ARNT: Aryl hydrocarbon nuclear translocator
bHLH: Basic helix loop helix
CYP1A1: Cytochrome P450 1A1
GSIS: Glucose stimulated insulin secretion
G6PC2: Glucose-6-phosphate catalytic subunit 2
HIF1α: Hypoxia inducible factor 1α
HRE: Hypoxia response element
IGRP: Islet-specific glucose-6-phosphatase catalytic subunit-related protein
MAFA: V-maf musculoaponeurotic fibrosarcoma oncogene homolog A
MNSOD: Manganese superoxide dismutase
PAS: Period/ARNT/Sim
POP: Persistent organic pollutant
SC-islets: Stem cell-derived islets
T2D: Type 2 diabetes
TCDD: 2,3,7,8 tetrachlorodibenzo-*p*-dioxin
XRE: Xenobiotic response element

## Notes

### Competing Interest Statement

The authors have declared no competing interest.

