## Supplemental Tables for "Crosstalk between the aryl hydrocarbon receptor and hypoxia inducible factor 1α pathways impairs downstream dioxin response in human islet models"

**Supplementary Table 1. SYBR qPCR primer sequences.**

| | |
|---|---|
| PPIA Forward<br>PPIA Reverse | AGCTCTGAGCACTGGAGAGA<br>GCCAGGACCTGTATGCTTTA |
| CYP1A1 Forward<br>CYP1A1 Reverse | ATCACAGACAGCCTCATTGAGC<br>AGATAGCAGTTGTGACTGTGTC |
| HMOX1 Forward<br>HMOX1 Reverse | TGACCCATGACACCAAGGAC<br>AGTGTAAGGACCCATCGGAGA |
| AHR Forward<br>AHR Reverse | TCCAAGCGGCATAGAGACCG<br>GCCTCCGTTTCTTTCAGTAGGG |
| AHRR Forward<br>AHRR Reverse | CTTAATGGCTTTGCTCTGGTCG<br>TGCATTACATCCGTCTGATGGA |
| VEGFA Forward<br>VEGFA Reverse | TTGCTCAGAGCGGAGAAAGC<br>GGTGAGAGATCTGGTTCCCG |
| ARNT Forward<br>ARNT Reverse | CGGAACAAGATGACAGCCTAC<br>ACAGAAAGCCATCTGCTGCC |
| MNSOD Forward<br>MNSOD Reverse | GGACAAACCTCAGCCCTAACGG<br>TTGGACACCAACAGATGCAGCC |
| SLC2A1 Forward<br>SLC2A1 Reverse | CTTCCAGTATGTGGAGCAACT<br>GAAGCGATCTCATCGAAGGT |
| MAFA Forward<br>MAFA Reverse | ATTCTGGAGAGCGAGAAGTGC<br>ATTTCTCCTTGTACAGGTCCCG |
| G6PC2 Forward<br>G6PC2 Reverse | GTCATCGACCTTACTGGTGGG<br>ACATACCAGACACAGGATGCG |

**Supplementary Table 2: F Statements for each human donor, by gene target.** Significance indicated by bolded text.

***CYP1A1***

| **Donor R425** | **F (DFn, DFd)** | **p value** |
|---|---|---|
| **Interaction effect** | **$F(1, 13) = 42.02$** | **$p < 0.001$** |
| **Hypoxia** | **$F(1, 13) = 70.59$** | **$p < 0.001$** |
| **TCDD** | **$F(1, 13) = 168.4$** | **$p < 0.001$** |
| **Donor R427** | | |
| Interaction effect | $F(1, 8) = 0.8521$ | $p > 0.05$ |
| Hypoxia | $F(1, 8) = 4.990$ | $p = 0.0560$ |
| **TCDD** | **$F(1, 8) = 55.77$** | **$p < 0.001$** |
| **Donor R430** | | |
| Interaction effect | $F(1, 15) = 4.322$ | $p = 0.0552$ |

| | | |
|---|---|---|
| Hypoxia | F (1, 15) = 1.813 | $p > 0.05$ |
| **TCDD** | **F (1, 15) = 55.98** | **$p < 0.001$** |
| **Donor R434** | | |
| Interaction effect | F (1, 15) = 0.4509 | $p > 0.05$ |
| Hypoxia | F (1, 15) = 0.3647 | $p > 0.05$ |
| **TCDD** | **F (1, 15) = 201.5** | **$p < 0.001$** |
| **Donor R444** | | |
| Interaction effect | F (1, 14) = 0.1532 | $p > 0.05$ |
| Hypoxia | F (1, 14) = 0.02673 | $p > 0.05$ |
| **TCDD** | **F (1, 14) = 292.3** | **$p < 0.001$** |
| **Donor R461** | | |
| **Interaction effect** | **F (1, 11) = 25.60** | **$p < 0.001$** |
| **Hypoxia** | **F (1, 11) = 13.45** | **$p < 0.01$** |
| **TCDD** | **F (1, 11) = 128.7** | **$p < 0.001$** |

*HMOX1*

| **Donor R425** | **F (DFn, DFd)** | **p value** |
|---|---|---|
| **Interaction effect** | **F (1, 13) = 8.536** | **$p < 0.05$** |
| **Hypoxia** | **F (1, 13) = 31.49** | **$p < 0.001$** |
| TCDD | F (1, 13) = 3.509 | $p > 0.05$ |
| **Donor R427** | | |
| Interaction effect | F (1, 8) = 0.07678 | $p > 0.05$ |
| **Hypoxia** | **F (1, 8) = 37.32** | **$p < 0.001$** |
| TCDD | F (1, 8) = 0.008461 | $p > 0.05$ |
| **Donor R430** | | |
| Interaction effect | F (1, 15) = 1.192 | $p > 0.05$ |
| **Hypoxia** | **F (1, 15) = 31.86** | **$p < 0.001$** |
| TCDD | F (1, 15) = 0.4044 | $p > 0.05$ |
| **Donor R434** | | |
| Interaction effect | F (1, 15) = 3.667 | $p > 0.05$ |
| **Hypoxia** | **F (1, 15) = 59.85** | **$p < 0.001$** |
| **TCDD** | **F (1, 15) = 6.891** | **$p < 0.05$** |
| **Donor R444** | | |
| Interaction effect | F (1, 14) = 3.410 | $p > 0.05$ |
| Hypoxia | F (1, 14) = 4.327 | $p = 0.0564$ |
| **TCDD** | **F (1, 14) = 4.863** | **$p < 0.05$** |
| **Donor R461** | | |
| Interaction effect | F (1, 11) = 0.3300 | $p > 0.05$ |
| **Hypoxia** | **F (1, 11) = 18.09** | **$p < 0.01$** |
| TCDD | F (1, 11) = 0.2796 | $p > 0.05$ |

*AHRR*

| **Donor R425** | **F (DFn, DFd)** | **p value** |
|---|---|---|
| **Interaction effect** | **F (1, 13) = 5.877** | **$p < 0.05$** |

| | | |
|---|---|---|
| Hypoxia | F (1, 13) = 0.5844 | p > 0.05 |
| **TCDD** | **F (1, 13) = 13.86** | **p < 0.01** |
| **Donor R427** | | |
| Interaction effect | F (1, 8) = 0.006853 | p > 0.05 |
| Hypoxia | F (1, 8) = 0.1562 | p > 0.05 |
| TCDD | F (1, 8) = 0.8378 | p > 0.05 |
| **Donor R430** | | |
| **Interaction effect** | **F (1, 15) = 6.636** | **p < 0.05** |
| Hypoxia | F (1, 15) = 0.07040 | p > 0.05 |
| **TCDD** | **F (1, 15) = 11.81** | **P < 0.01** |
| **Donor R434** | | |
| Interaction effect | F (1, 15) = 0.7542 | p > 0.05 |
| Hypoxia | F (1, 15) = 2.271 | p > 0.05 |
| **TCDD** | **F (1, 15) = 9.734** | **p < 0.01** |
| **Donor R444** | | |
| Interaction effect | F (1, 15) = 1.633 | p > 0.05 |
| Hypoxia | F (1, 15) = 1.922 | p > 0.05 |
| **TCDD** | **F (1, 15) = 25.44** | **p < 0.001** |
| **Donor R461** | | |
| Interaction effect | F (1, 11) = 0.06245 | p > 0.05 |
| Hypoxia | F (1, 11) = 1.348 | p > 0.05 |
| **TCDD** | **F (1, 11) = 37.26** | **p < 0.001** |

***ARNT***

| **Donor R425** | **F (DFn, DFd)** | **p value** |
|---|---|---|
| Interaction effect | F (1, 12) = 1.113 | p > 0.05 |
| Hypoxia | F (1, 12) = 0.5568 | p > 0.05 |
| TCDD | F (1, 12) = 2.086 | p > 0.05 |
| **Donor R427** | | |
| Interaction effect | F (1, 8) = 0.5742 | p > 0.05 |
| Hypoxia | F (1, 8) = 2.132 | p > 0.05 |
| TCDD | F (1, 8) = 0.2949 | p > 0.05 |
| **Donor R430** | | |
| **Interaction effect** | **F (1, 15) = 4.830** | **p < 0.05** |
| Hypoxia | F (1, 15) = 0.1023 | p > 0.05 |
| TCDD | F (1, 15) = 1.761 | p > 0.05 |
| **Donor R434** | | |
| Interaction effect | F (1, 15) = 0.1312 | p > 0.05 |
| **Hypoxia** | **F (1, 15) = 5.671** | **p < 0.05** |
| **TCDD** | **F (1, 15) = 6.179** | **p < 0.05** |
| **Donor R444** | | |
| Interaction effect | F (1, 15) = 1.894 | p > 0.05 |
| Hypoxia | F (1, 15) = 4.512 | p = 0.0507 |
| **TCDD** | **F (1, 15) = 26.73** | **p < 0.001** |
| **Donor R461** | | |

| | | |
|---|---|---|
| Interaction effect | F (1, 11) = 0.001704 | p > 0.05 |
| Hypoxia | F (1, 11) = 2.559 | p > 0.05 |
| TCDD | F (1, 11) = 1.678 | p > 0.05 |

***G6PC2***

| **Donor R425** | **F (DFn, DFd)** | **P value** |
|---|---|---|
| Interaction effect | F (1, 13) = 0.8402 | p > 0.05 |
| **Hypoxia** | **F (1, 13) = 41.18** | **p < 0.001** |
| TCDD | F (1, 13) = 0.007048 | p > 0.05 |
| **Donor R427** | | |
| Interaction effect | F (1, 8) = 1.915 | p > 0.05 |
| **Hypoxia** | **F (1, 8) = 6.509** | **P < 0.05** |
| TCDD | F (1, 8) = 1.863 | p > 0.05 |
| **Donor R430** | | |
| Interaction effect | F (1, 15) = 1.158 | p > 0.05 |
| **Hypoxia** | **F (1, 15) = 104.5** | **p < 0.001** |
| TCDD | F (1, 15) = 2.098 | p > 0.05 |
| **Donor R434** | | |
| Interaction effect | F (1, 15) = 1.431 | p > 0.05 |
| **Hypoxia** | **F (1, 15) = 11.72** | **p < 0.01** |
| **TCDD** | **F (1, 15) = 4.945** | **p < 0.05** |
| **Donor R444** | | |
| Interaction effect | F (1, 15) = 0.002809 | p > 0.05 |
| Hypoxia | F (1, 15) = 1.086 | p > 0.05 |
| **TCDD** | **F (1, 15) = 11.99** | **p < 0.01** |
| **Donor R461** | | |
| Interaction effect | F (1, 11) = 0.08265 | p > 0.05 |
| **Hypoxia** | **F (1, 11) = 76.44** | **p < 0.001** |
| TCDD | F (1, 11) = 0.1775 | p > 0.05 |
